# xTrimoDock: Rigid Protein Docking via Cross-Modal Representation Learning and Spectral Algorithm

**DOI:** 10.1101/2023.02.06.527251

**Authors:** Yujie Luo, Shaochuan Li, Yiwu Sun, Ruijia Wang, Tingting Tang, Beiqi Hongdu, Xingyi Cheng, Chuan Shi, Hui Li, Le Song

**Author notes:** Equal contribution (Yujie Luo, Shaochuan Li).

## Abstract

Protein-protein interactions are the basis for the formation of protein complexes which are essential for almost all cellular processes. Knowledge of the structures of protein complexes is of major importance for understanding the biological function of these protein-protein interactions and designing protein drugs. Here we address the problem of rigid protein docking which assumes no deformation of the involved proteins during interactions. We develop a method called, xTrimoDock, which leverages a cross-modal representation learning to predict the protein distance map, and then uses a spectral initialization and gradient descent to obtain the roto-translation transformation for docking. We show that, on antibody heavy-chain and light-chain docking, and antibody-antigen docking, xTrimoDock consistently outperforms the state-of-the-art such as AlphaFold-Multimer and HDock, and can lead to as much as a 10% improvement in DockQ metric. xTrimoDock has been applied as a useful tool in protein drug design at BioMap.

## 1 Introduction

Protein-protein interactions (PPIs) are the basis of many fundamental cellular processes, including inhibition or activation of enzymes, cellular signaling, and recognition of antigens by the adaptive immune system. High-resolution structural characterization of these interactions provides insights to understand their biological function and facilitates the discovery of protein-based drugs. However, the experimental gold standards for determining the structure of protein complexes, such as X-ray crystallography and cryo-electron microscopy (cryo-EM), are costly and time-consuming.

Computational protein docking offers an alternative way to predict the three-dimensional structures of protein complexes from the structures of the two or more proteins in their unbound states (Venkatraman et al., 2009; Weng et al., 2019). Here, we focus on the common problem of rigid protein docking where no deformations of the involved individual proteins occur within the binding process. Therefore, an appropriate roto-translation transformation needs to be estimated to place the ligand at the right location and orientation with respect to the receptor, as shown in Figure 1.

**Figure 1:**
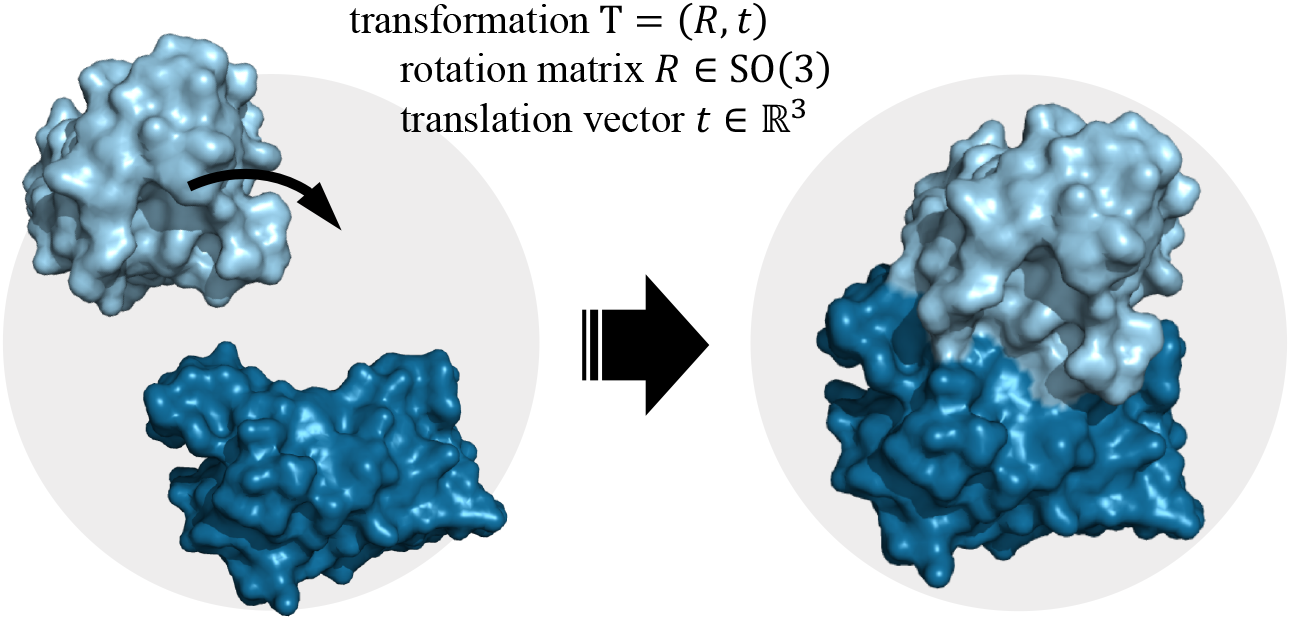
Surface views of rigid protein docking. Keeping one protein at a fixed location, and a roto-translation transformation *T* = (*R, t*) is predicted to place another protein at the right location and orientation.

Classical docking software (Chen et al., 2003; De Vries et al., 2010; Kozakov et al., 2017; Yan et al., 2020; Eismann et al., 2021) largely follows a costly three-step framework: i) samples a large number of candidate structures, ii) leverages a scoring function to rank them, and iii) refines the top structures based on an energy model. Due to a large number of generated samples and score evaluation, these docking algorithms are very slow. Furthermore, the scoring functions of these docking algorithms are based on simple statistics computed from protein complexes in the PDB database, making them very sensitive to noise. If the involved individual protein structures are inaccurate because, for instance, they are obtained from an inaccurate structure prediction algorithm, then the performance of these docking algorithms may be severely impacted.

Recently, a deep learning model called EquiDock (Ganea et al., 2022) was proposed for rigid protein docking and results in a great reduction in computational time compared to classical docking algorithms. Although EquiDock makes good use of the involved individual protein structures and the non-deformable physical process of rigid docking, it cannot incorporate useful information hidden in protein sequences and protein sequence databases, such as multiple sequence alignments (MSA) from a query protein to a protein sequence database. Recently, it has been shown that MSA plays a critical role in improving deep learning based protein structure prediction and protein complex structure prediction (Jumper et al., 2021; Evans et al., 2021). Prominently, AlphaFold-Multimer (Evans et al., 2021) infers a protein complex structure from protein sequences and MSA alone without using any individual protein structures and achieves state-of-the-art performance. However, AlphaFold-Multimer does not respect the physical process of rigid docking, i.e., no protein deformation, and it can lead to unnatural protein complex structures. In sum, existing deep learning-based methods fall short in either making full use of multi-modal information in proteins (sequences and structures), or the physics of rigid docking.

### Contribution

To address the above limitations, we propose a deep learning method called xTrimoDock, which is fast and incorporates both multi-modal information and physics. More specifically, xTrimoDock leverages Evoformer architecture (Jumper et al., 2021) to learn cross-modal representation from both protein sequences, in particular MSAs, and single protein structures, and then the representation is used to predict the cross-protein distance map.

Then the roto-translation transformation is learned via the minimization of the difference between the predicted distance map and the actual distance map from docking. This is a nonconvex optimization problem which is not easily solved. Thus, we derive a spectral algorithm for initialization followed by further refinement with gradient descents. We conduct experiments on variable region of antibody heavy-chain and light-chain docking, and antibody-antigen docking, both of which are important for antibody drug design. xTrimoDock achieved a 10% improvement in DockQ metric compared to AlphaFold-Multimer, showing the promise of our proposed xTrimoDock as a tool for antibody drug design.

## 2 Backgrounds and Related Works

We will first provide some details on backgrounds and relevant works below.

### 2.1 Antibody and antibody-antigen docking

Antibodies are proteins produced by the immune system and play a critical role in recognizing and neutralizing foreign invaders such as viruses and bacteria. They are made up of four polypeptide chains, two identical heavy chains and two identical light chains, that are connected together through disulfide bonds to form a Y-shaped structure. The part of the antibody that interacts with antigens is called the antigen-binding site, and it is located at the tip of the two arms of the Y-shaped structure. The fragment of variable region (Fv region) is made up of two variable regions, VH and VL, containing complete antigen-binding region. Both VH and VL contain three hypervariable regions, the complementarity-determining regions (CDRs), which serve as key structural parts of the antibody-antigen binding site in antibodies. Antigen-antibody docking refers to the process by which an antibody and an antigen bind together. The antibody recognizes the antigen in a specific position, or more specifically they form an antibody-antigen complex where the antibody docks to the antigen with a specific roto-translation transform. In the Ab-Ag (antibody-antigen) complex, the antibody contains a paratope (analogous to a key) that is specific for one particular epitope (analogous to a lock) on an antigen, allowing these two structures to bind together with precision. As in Figure 2, since an antibody has a *H*_2_*L*_2_ symmetric sequence feature, and the antibody-antigen interaction occurs at an Fv region of the antibody, we will focus on the structure of an Fv region for the antibody, and the complex structure of an Fv with the antigen. For convenience, we will refer to the Fv-antigen complex as the antibody-antigen complex in the remainder of the paper.

**Figure 2:**
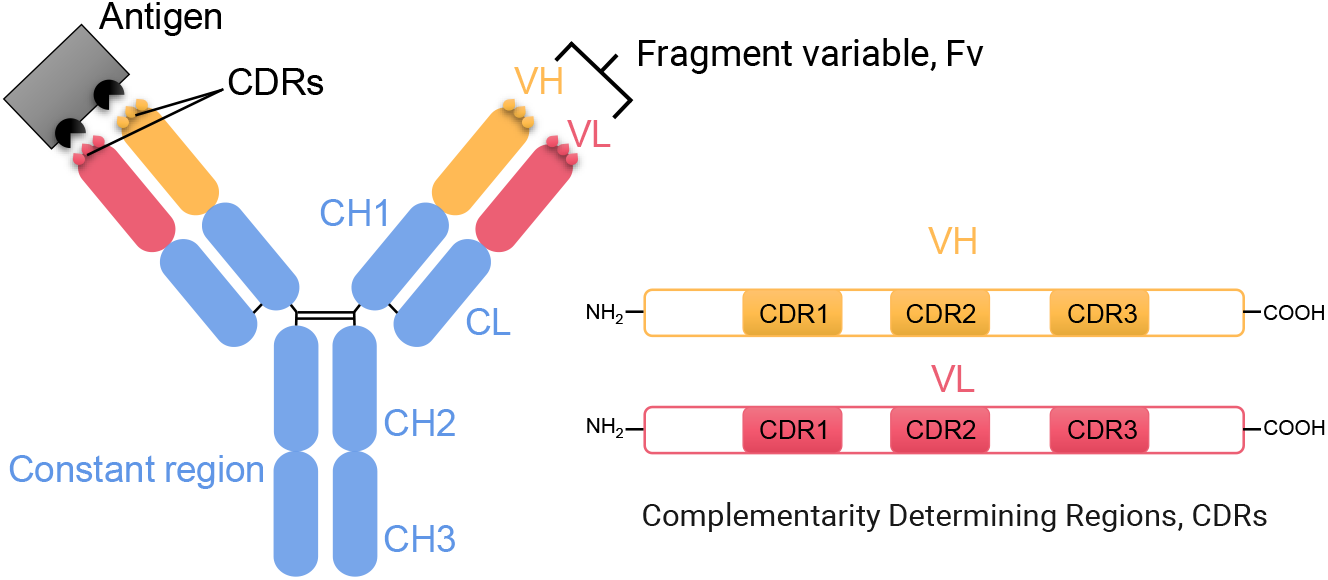
The schematic representation of an IgG antibody and the antibody-antigen interaction. The fragment of variable region (Fv region) of heavy chain and light chain is colored orange and red, respectively. Constant regions including CL, CH1, CH2 and CH3 are colored blue. Antigen is colored gray. Both VH and VL contain three highly variable complementarity-determining regions (CDRs), which are more important than other regions for antibody-antigen docking.

### 2.2 Protein structure prediction

Very often, designed antibody drugs or the antigen may not have 3D structures. Their structures need to be predicted from protein sequences before docking algorithms can be applied. Protein structure prediction is an important research problem that gets the 3D coordinates from a given amino acid sequence. In recent years, deep learning methods have been widely used in protein structure prediction, such as AlphaFold (Senior et al., 2019), AlphaFold2 (Jumper et al., 2021), Openfold (Ahdritz et al., 2022) and RoseTTAFold (Baek et al., 2021). Using the co-evolution information from Multiple Sequence Alignments (MSA), these methods achieve high accuracy on both backbone and side-chain predictions. Other structure prediction methods, such as OmegaFold (Wu et al., 2022), Helixfold-single (Fang et al., 2022), ESM-Fold (Lin et al., 2022), employ large-scale protein language models to replace computationally extensive MSA searching and achieve comparable performance with AlphaFold2. Models designed specifically for antibody structure prediction, such as DeepAb (Ruffolo et al., 2022), ABlooper (Abanades et al., 2022a), IgFold (Ruffolo and Gray, 2022), xTrimoABFold (Wang et al., 2022), ImmuneBuilder (Abanades et al., 2022b) have recently been developed. In particular, xTrimoABFold uses a pretrained antibody language model and crossmodal template searching algorithm to obtain the most accurate heavy or light chain structure of an antibody.

Antibody-antigen docking is a computational method used to predict the binding between an antibody and its antigen. The algorithm uses structural and chemical information about the antibody and antigen to determine the most stable and energetically favorable binding mode.

Computational docking methods can be generally divided into classical docking software and deep learning-based methods (Biesiada et al., 2011; Torchala et al., 2013; Vakser, 2014; Schindler et al., 2017; Weng et al., 2019; Yan et al., 2020; Sunny and Jayaraj, 2021; Christoffer et al., 2021). Specifically, classical docking software is typically very slow and follows a three-step framework of candidate sampling, ranking (Launay et al., 2020; Eismann et al., 2021), and refinement (Verburgt and Kihara, 2022). Since deep learning has made a big impact on structural biology (Laine et al., 2021), AlphaFold2 (Jumper et al., 2021) and RoseTTAFold (Baek et al., 2021) have also been utilized to predict protein complex structures by adding a sequence linker between individual protein sequences.

Recently, AlphaFold-Multimer (Evans et al., 2021) extends AlphaFold2 to fold and dock two proteins by explicitly linking the MSAs from the two proteins and training the model with protein complex structures. However, AlphaFold-Multimer does not exploit the physics of rigid docking. EquiDock (Ganea et al., 2022) is tailored for effective rigid docking, but can not incorporate information from MSA modality.

## 3 Method

In this section, we propose a deep learning method, xTrimoDock, for antibody heavy chain and light chain docking, and for antibody-antigen docking. The design of xTrimoDock leverages multi-modal information, including multiple sequence alignments (MSAs) and protein structures, as well as incorporating physics. Briefly, as shown in Figure 3, xTrimoDock adopts protein primary structures as input and has four essential components before delivering the final complex 3D structure. The four components are 1) single-chain structure determination (section 3.1), 2) the distance map prediction model to obtain multi-chain relationships (section 3.2), 3) the fast spectral estimation method to achieve coarse rotation and translation according to the derived closed-form solution (section 3.3), and 4) the gradient refinement method to improve rotation and translation (section 3.4), respectively. Since xTrimoDock does not need to sample a large number of candidate decoys and uses deep learning approach to learn rotation and translation directly, it is very fast compared to classical docking methods. In the following subsections, we introduce the details for each component step by step.

**Figure 3:**
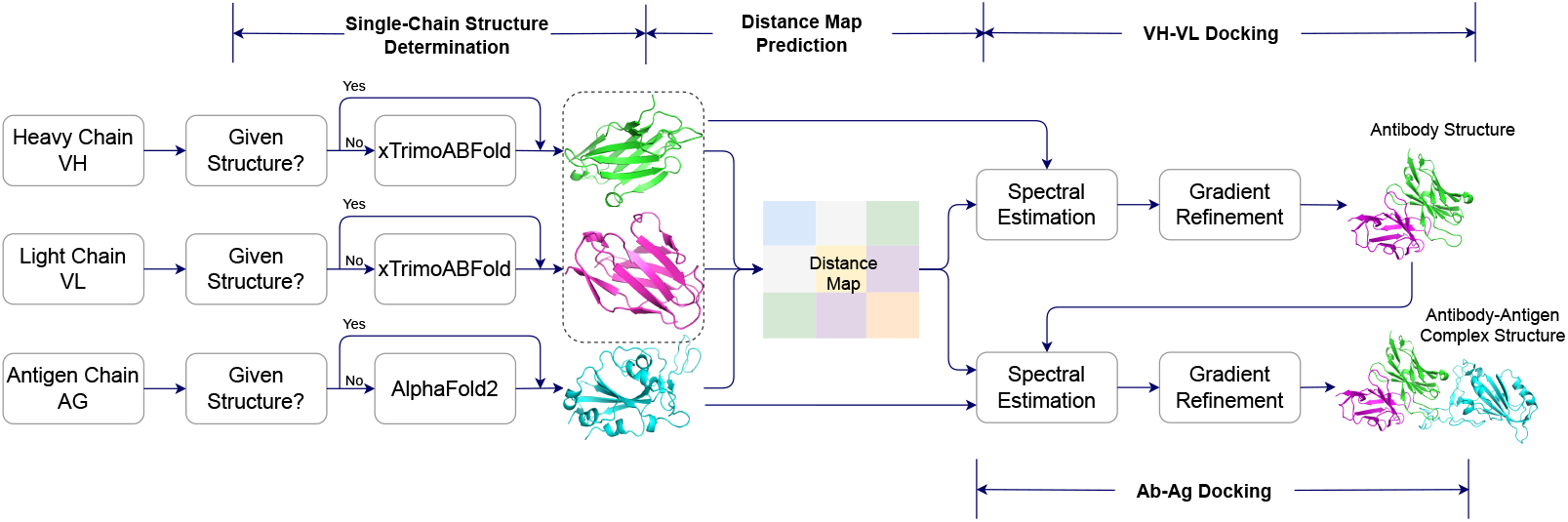
An illustration of the xTrimoDock framework. The xTrimoDock pipeline consists of four essential components: single-chain structure determination, distance map prediction model, spectral estimation model, and gradient refinement model.

### 3.1 Single-chain structure determination

The basic input for rigid docking is the single-chain’s rigid structure. However, there is no available rigid structure in the actual scenario, therefore, we applied the current state-of-the-art protein folding prediction model to generate the rigid structure. We employ xTrimoABFold (Wang et al., 2022) to predict structures for antibody heavy chain and light chain, and employ AlphaFold2 (Jumper et al., 2021) to predict a structure for the antigen chain if there are no available structures experimentally measured by X-ray crystallography, cryo-EM, NMR spectroscopy, dual polarization interferometry, or other methods.

Given single-chain structures, we design a distance map prediction model to estimate pairwise distance of residues within multiple chains. The distance map prediction model contains three components as illustrated in Figure 4: 1) preparing model inputs from multiple modalities including primary sequences, single-chain structures and evolutionary information (paired MSAs), 2) evoformer module to learn pair representation and coevolution representation, 3) loss function containing both supervised distogram loss and self-supervised masked language model loss.

**Figure 4:**
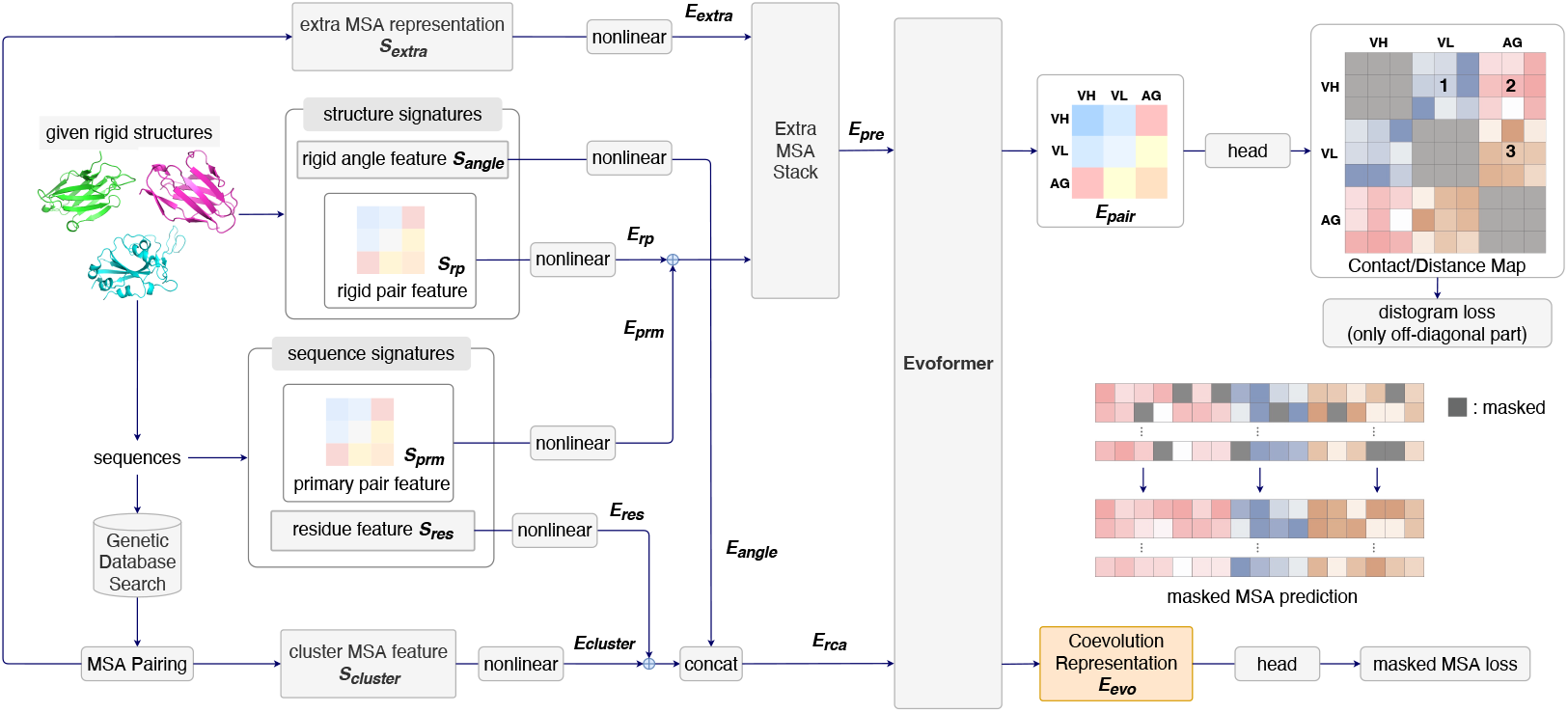
Cross-modal representation learning framework for distance map prediction on multiple proteins. The input contains three modalities: rigid structures, protein sequences and multiple sequence alignments (MSAs) searched from genetic databases. Because the rigid structures are given and distance map is symmetrical, we need to predict upper off-diagonal distance maps only, i.e., the distance maps 1, 2, 3 highlighted above.

### 3.2 Distance map prediction via cross-modal representation learning

#### 3.2.1 Multimodal Model Inputs

We will first process the three modes of inputs, i.e. structure, MSA and sequence, into their corresponding feature signatures as follows. Because of the space limitation, we present brief introduction for each mode of feature in this section and give further details in appendix D.

##### Structure Signatures

We derive two types of features from single-chain structures: *rigid angle feature* 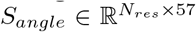 and *rigid pair feature* 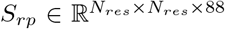. The *rigid angle feature* provides the orientation and position for each amino acid, and the *rigid pair feature* provides the distance information between two amino acids.

##### MSA Signatures

MSAs are crucial tools for comparing and analyzing the evolutionary relationships among different protein sequences. Cross-chain coevolutionary information obtained from MSAs is a useful feature in distance predictions. We employ the heuristic approach in (Evans et al., 2021) to pair sequences between per-chain MSAs. Specifically, the per-chain MSA sequences are grouped by species, where the species labels derived from UniProt’s id mapping ^3^. Then the grouped sequences are paired within a specific species. We concatenate the sequences if there is only one sequence per species. Otherwise, we match the chain MSAs by minimizing the base-pair distance between chains for prokaryotic species and match the chain MSAs by ordering them by sequence identity to the target sequence for eukaryotic species (Zhou et al., 2018). To reduce the computational and memory cost, we select *N*_*cluster*_ sequences randomly as MSA cluster centers, where we set the first cluster center always be the primary protein sequence. The remaining sequences are assigned to their closest cluster by Hamming distance. Also, we select *N*_*extra*_ sequences randomly without replacement from sequences that have not been selected as cluster centers. Finally, we derive two types of MSA features: *cluster MSA feature* 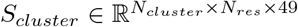 and *extra MSA feature* 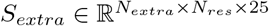.

##### Sequence Signatures

We derive two types of feature from primary sequence: *residue feature* 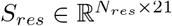 and *primary pair feature* 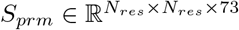.

#### 3.2.2 Learning Representation via Evoformer

Figure 4 introduces the neural architecture for our cross-modal distance prediction model. First, we use three fully connected neural networks (FCNNs) to learn representations from *residue feature S*_*res*_, *rigid angle feature S*_*angle*_ and *cluster MSA feature S*_*cluster*_ respectively, which is formulated by

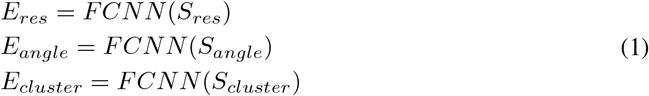

Then we add the learned *E*_*res*_ and *E*_*cluster*_ before concatenating *E*_*angle*_, which is formulated by

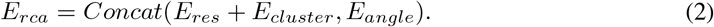

Similarly, we use another fully connected neural networks (FCNNs) to learn representations from *primary pair feature S*_*prm*_, *rigid pair feature S*_*rp*_ and *extra MSA feature S*_*extra*_.

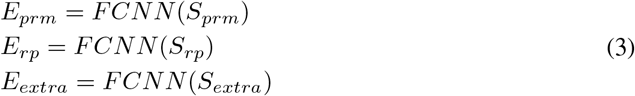

Then we add *E*_*prm*_ to *E*_*rp*_, and the additive sum together with *E*_*extra*_ are fed into an Extra MSA Stack network introduced in (Jumper et al., 2021).The above process is formulated by

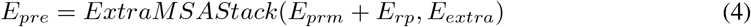

We employ evoformer (Jumper et al., 2021) to learn pair representation *E*_*pair*_ and coevolution representation *E*_*evo*_ from the learned *E*_*pre*_ and *E*_*rca*_.

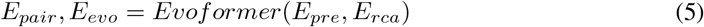

Finally, we use two heads (FCNNs) separately to obtain the final pair representation and coevolution representation.

#### 3.2.3 Loss Functions

Define *d*_*ij*_ ∈ ℝ as the ground truth distance map. Define neural network function *f*_*θ*_(*x*) ∈ ℝ parameterized by *θ* be the predicted distance map. We formulate distance prediction model by

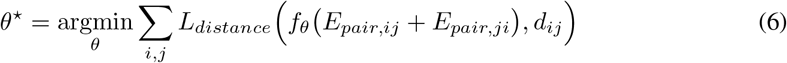

Typically, the above function could be optimized by minimizing the standard mean squared error (MSE), however, *d*_*ij*_ is naturally noisy because of the experimentally-determined 3D protein structures. According to the practices in the field of protein structure predictions (Wu and Zhang, 2008; Jumper et al., 2021; Evans et al., 2021), we use discretized distance map (distogram, see Figure 4) to replace the exact distance map. Specifically, the distance between two residues is discretized into 64 bins which cover from 2 *Å* to 22 *Å* with the middle of the bins denoted as {*δ*_1_, *δ*_2_, …, *δ*_64_}. The task of distance map prediction is then turned into a classification problem. Define 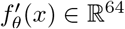 as the neural network to predict discretized distance. Define 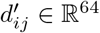 as the one-hot encoding of actual discretized distance. Define 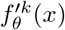 and 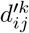 as the *k*-th value in 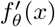 and 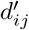. We calculate cross-entropy loss between 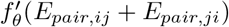 and 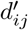.

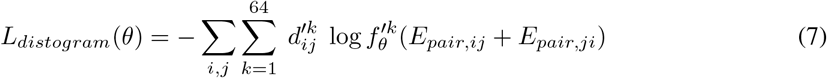

To further learn coevolutionary information from pairing cross-chain MSAs, we design a neural network *g*_*η*_ parameterized by *η* to learn the masked MSA loss, which is formulated by

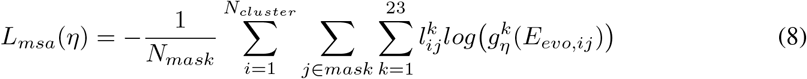

where *N*_*mask*_ refers the the number of masked tokens, 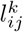 refers to the *k*-th ground-truth value encoded by one hot, 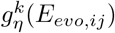 refers to the *k*-th predicted class probability. We introduce the mask policy for MSA in appendix B.

Our final objective function is formulated by

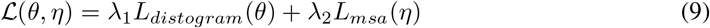

where *λ*_1_ and *λ*_2_ are two weights for distogram loss and masked MSA loss.

Equation 9 is excepted to be trained on protein complexes from the Protein Data Bank (Berman, 2008).

The expected distance between residue *i* and *j* is defined by 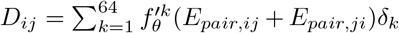.

### 3.3 Spectral Estimation with Rigid Structures

With information on the rigid structure and distance map, we initiate the docking procedure. Our docking algorithm is trying to learn a rotation and translation so that the distance information between two rigid structures is similar to the input distance map, i.e., the objective function ℒ (*R, t*) is defined as follows:

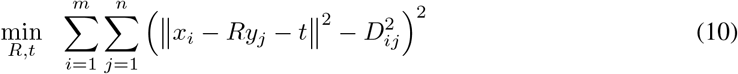

where *x*_*i*_ ∈ ℝ^3×1^ and *y*_*j*_ ∈ ℝ^3×1^ is the given initial beta carbon coordinate of *i*-th and *j*-th residue extracted from the given rigid structure, ℝ ∈ ℝ^3×3^ is the rotation matrix, *t* ∈ ℝ^3^ is the translation vector, and *D*_*ij*_ is the expected distance predicted by *f*_*θ*_, *m* and *n* refers to the number of residues in two rigid structures.

However, the problem in Equation (10) is nonconvex, and gradient descent algorithm will easily get stuck in local minima as illustrated by figure 5. To address this problem, we develop a spectral initialization approach, leveraging on a sufficient condition for the optimal solution of Equation (10).

**Figure 5:**
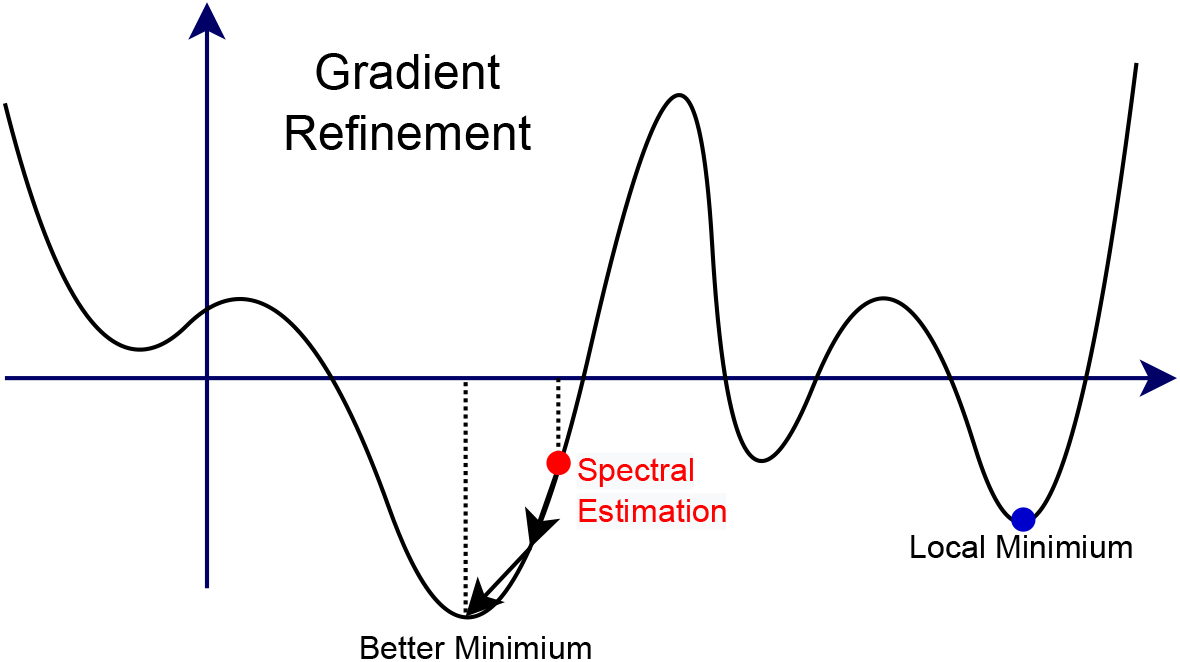
Gradient refinement from spectral estimation.

To simplify Equation (10), define *ŷ*_*i*_ = *Ry*_*i*_+*t, X* = [*x*_1_, *x*_2_, …, *x*_*m*_] ∈ *ℝ*^3*×m*^, *Y* = [*y*_1_, *y*_2_, …, *y*_*n*_] ∈ ℝ^3*×n*^, *Ŷ* = *RY* + *t* = [*ŷ*_1_, *ŷ*_2_, …, *ŷ*_*n*_] ∈ *ℝ*^3*×n*^,

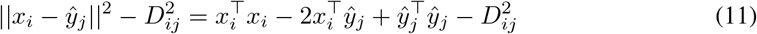

Furthermore, we define *A, B, C, D* ∈ *ℝ*^*m×n*^ by

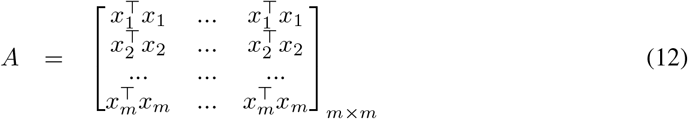

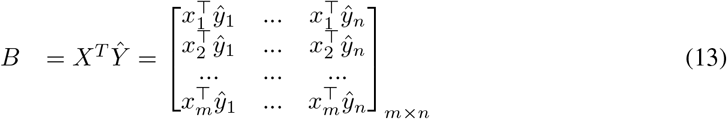

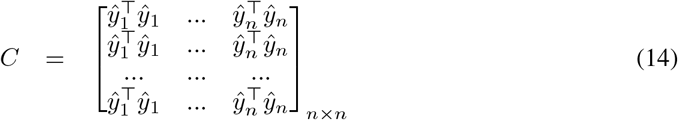

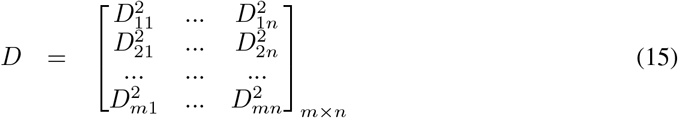

If the predicted distance map *D* is equal to its ground truth, we can obtain the closed-form *R* and *t* for Equation (10), satisfying the following condition: ∀*i, j, s*.*t*., ||*x*_*i*_ − *ŷ*_*j*_|| = *D*_*ij*_, *i*.*e*.,

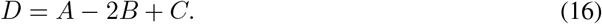

Further define two centering matrices

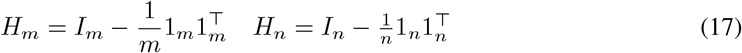

respectively, and we observe that

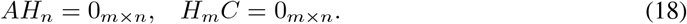

Then applying *H*_*m*_ and *H*_*n*_ to the left side and right side of Equation (16) respectively, and we obtain

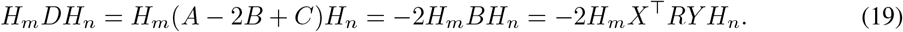

We perform thin singular value decompositions (SVD) for *H*_*m*_*X*^⊤^ and *Y H*_*n*_ respectively,

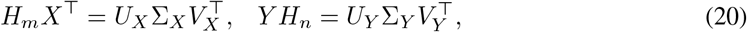

where both ∑_*X*_ and ∑_*y*_ are 3 by 3 diagonal matrices. Then the rotation matrix *R* can be solved as

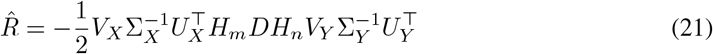

After we solve *R*, we can further solve for *t* using

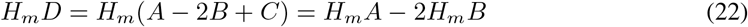

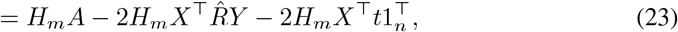

that is

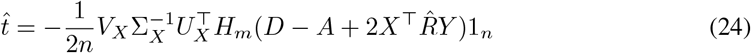

### 3.4 Gradient Refinement

Even if we can calculate the rotation and translation matrices from the rigid structure and the distance map in previous section, it is not sufficient because of the noisy predicted distance. To obtain the more accurate rotation and translation matrices, we further employ the gradient descent method to refine the rotation matrix 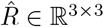 and translation matrix 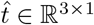. Algorithm 1 gives the pseudo code for gradient refinement.

#### Rotation Reparameterization

We need to reparameterize rotation matrix 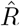 before conducting gradient descent. An orthogonal rotation matrix *R* is determined by a unit quaternion ***q*** = *a* + *b****i*** + *c****j*** + *d****k***, *i*.*e*., we can calculate *R* by

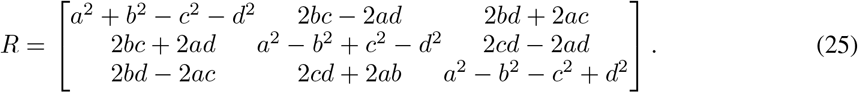

##### Algo 1

Gradient Refinement

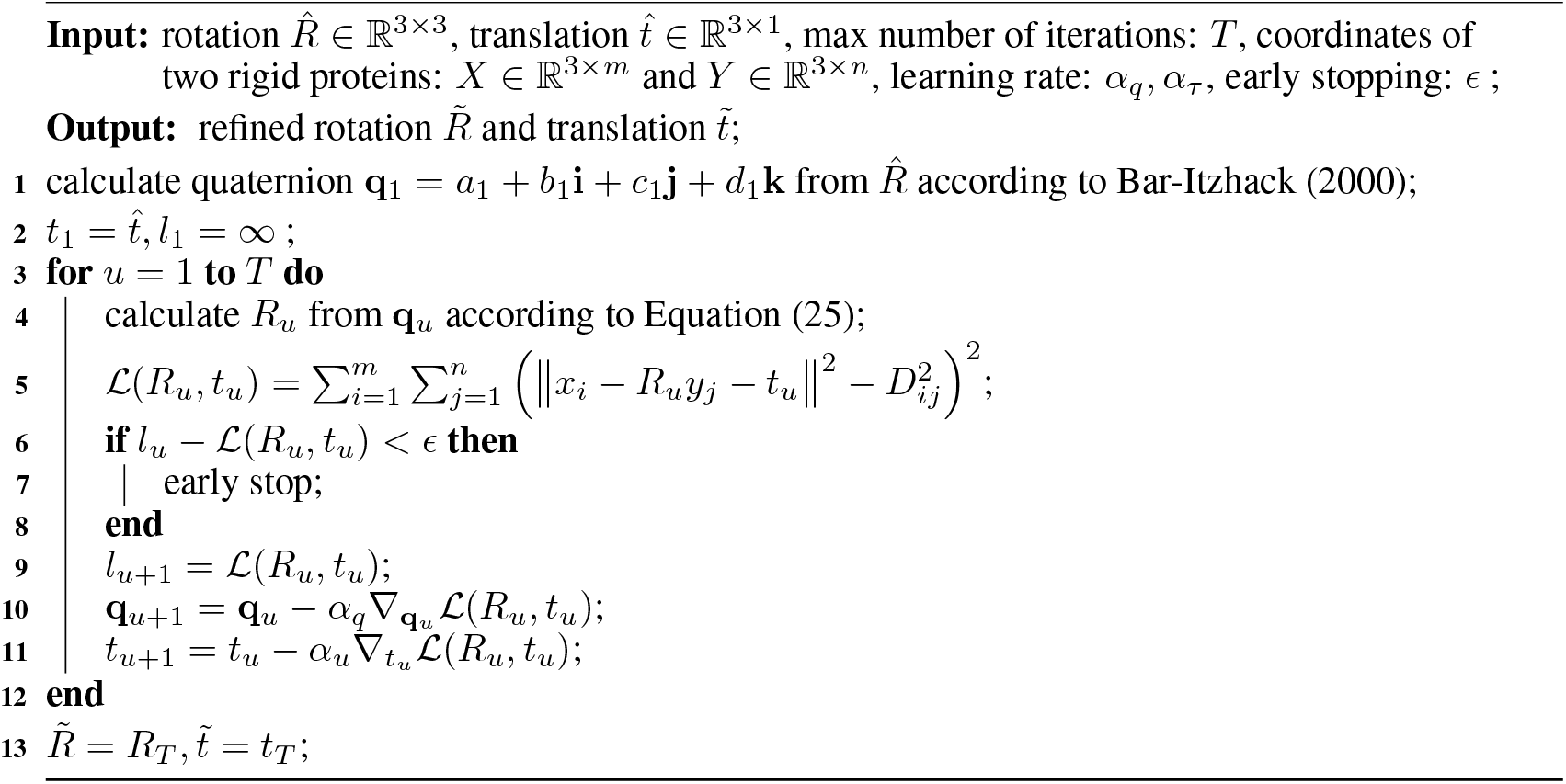

Given the estimated rotation 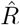 from previous section, we can obtain the quaternion ***q*** = *a*+*b****i***+*c****j***+*d****k*** corresponding to the rotation matrix closest to the given matrix 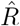 using a method by Bar-Itzhack (2000). Specifically, we construct a symmetric 4 × 4 matrix *K*

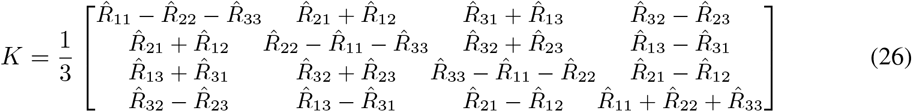

and can find the eigenvector, (*b, c, d, a*), corresponding to its largest magnitude eigenvalue. Then this obtained quaternion ***q*** = *a* + *b****i*** + *c****j*** + *d****k*** can be used to construct a rotation matrix *R* as in Equation (25) corresponding to the rotation matrix closest to the given matrix 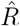.

#### Refinement by Gradient Descent

After reparameterizing rotation 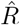 by its corresponding quaternion, we conduct gradient descent on quaternion and translation matrix 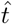 to refine the solution. After the gradient refinement, we obtain the optimized 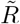 and 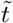 for rigid docking.

As illustrated in Figure 5, spectral estimation provides a good initialization for gradient refinement which helps gradient descent to avoid getting stuck in a local minimum and accelerates the whole process. Both spectral estimation and gradient refinement are indispensable parts of the algorithm. Furthermore, spectral estimation ensures the stability of our algorithm and gradient refinement eliminates the effects of noisy distance maps.

## 4 Experiment

In the section, we evaluate xTrimoDock on recently released antibody-antigen complex and compare xTrimoDock with the state-of-the-art docking methods.

### 4.1 Experimental Setup

#### Datasets

For distance map training, our training set is created from the Protein Data Bank (PDB) (Berman, 2008), which contains massive experimentally-determined 3D protein structures. The training set consists of 5,804 complexes, each containing proteins with at least 30 residues. The complexes in training set contain two or more chains including at least one antibody chain and the average chain number of the train set is 3.1. All complexes in training set are released before January 12, 2022.

For the experimental test, we create an antibody-antigen dataset from PDB, consisting of 68 antibody-antigen complexes as listed in appendix A. To make a fair comparison, all the curated antibody-antigen complexes are released after October 2022 and have not been used to train EquiDock, AlphaFold-Multimer and xTrimoDock. Specifically, the antibody-antigen complex is composed of 3 chains, including one light chain, one heavy chain from the antibody, and one chain from the antigen. Then we extract the antibody from the complexes to form an antibody dataset. The antibody complexes are composed of 2 chains, including a light chain and a heavy chain. These two chains form the basic functional part of an antibody, often referred as fragment variable region (Fv), which is the key part for antigen binding. The statistics of the datasets are summarized in Table 1.

**Table 1:** Statistics of datasets. VH-VL refers to the dataset for antibody VH and VL docking. Ab-Ag refers to the dataset for antibody-antigen docking.

|  | Residues per chain |  |  | Atoms per chain |  |  |
| --- | --- | --- | --- | --- | --- | --- |
|  | MIN | MAX | MEAN | MIN | MAX | MEAN |
| Train set | 30 | 1425 | 189 | 213 | 16951 | 1846 |
| Test set (VH-VL) | 103 | 132 | 115 | 730 | 1979 | 952 |
| Test set (Ab-Ag) | 30 | 430 | 143 | 248 | 3270 | 1177 |

#### Baselines

To evaluate the effectiveness of our proposed model, we compare xTrimoDock with two categories of methods, including three classical docking softwares: ZDOCK (Chen et al., 2003), ClusPro (Kozakov et al., 2017), HDOCK (Yan et al., 2020) and two deep learning methods, AlphaFold-Multimer (Evans et al., 2021), EquiDock (Ganea et al., 2021). ZDOCK, ClusPro and HDock provide local packages for convenient offline evaluation or webservers for online submissions. For AlphaFold-Multimer ^4^ and EquiDock ^5^, we use their released models with default settings on GitHub.

#### Metrics

To evaluate the quality of predictions, we use Root Mean Squared Deviation (RMSD), Template-Modeling Score (TM-Score (Zhang and Skolnick, 2004)) and DockQ (Basu and Wallner, 2016) as evaluation metrics, which are widely used in protein docking (Jumper et al., 2021; Evans et al., 2021). DockQ is a weighted average of three terms including contact accuracy, interface RMSD, and ligand RMSD. It is a score in the range [0, 1] that can be used to measure the quality of the interface. Interfaces with scores greater than 0.23 are considered as successfully predicting the interfaces, greater than 0.49 and less than 0.8 are considered as medium quality, and greater than 0.8 are considered as high quality(Basu and Wallner, 2016). To further evaluate the docking quality, we define the Medium-High Quality (MHQ) metric, referring to the percentage of cases whose DockQ are greater than 0.49. We compute RMSD, TM-Score in USalign (Zhang et al., 2022) and compute DockQ in the public tool ^6^.

#### Implementation

For most real-world applications of antibody-antigen docking, single-chain’s rigid structures are not available because of the expensive experimentally structure determination. The stat-of-the-art computational protein folding models provide alternative solutions and provide protein structures from single-chain primary sequences with atomic accuracy. In this paper, we predict the structures of antibody’s light chains and heavy chains by xTrimoABFold (Wang et al., 2022) and antigen’s structures by AlphaFold2 (Jumper et al., 2021). To eliminate the effects of MSAs, we use the same MSAs for antigen sequences on AlphaFold2 and AlphaFold-Multimer. Baseline models, except AlphaFold-Multimer, are designed for binary complex setting, so that the heavy chain and light chain of the antibody are docked prior to antibody-antigen docking.

#### Reproducibility

In terms of xTrimoDock, for distance map training, the MSA clustering approach is adapted from AlphaFold2 (Jumper et al., 2021), which uses JackHMMER (Johnson et al., 2010) and HHBlits (Remmert et al., 2012). We select *N*_*cluster*_ = 252 and *N*_*extra*_ = 1024. The outputs of MSAs are deduplicated, stacked, and cropped to 49 dimensions while the extra MSAs outputs are cropped to 25 dimensions.The random sequence crop size is set to 412 and the embedding size of pair representation and coevolution representation is set to 128 and 256 respectively. As for objective function in Equation 9, we set *λ*_1_=0.3 and *λ*_2_=2. The evoformer contains 48 layers. The distogram prediction head 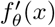 contains 1 fully connected layer. The mask MSA prediction head *g*_*η*_(*x*) contains 1 fully connected layer, too. We use Adam optimizer with *β*_1_=0.9, *β*_2_=0.999. The learning rate warms up from 0 to 1e-4 lasting for 1000 steps. After that, the learning rate decays by 5% every 5000 steps. We train our model on 8 Nvidia A100 GPUs with a batch size of 1 on each GPU. During gradient refinement phase, we use Adam optimizer with learning rate *α*_*q*_ = *α*_*τ*_ =1e-1 and *β*_1_ = 0.9, *β*_2_ = 0.999 and gradient refinement step *T* = 10000 with early stopping *ϵ* = 1*e*− 4 in Algorithm 1.

### 4.2 Results and Analysis

#### Antibody heavy chain and light chain docking

The performance comparison of xTrimoDock with other baselines is shown in Table 3 and Figure 6. Both xTrimoDock and AlphaFold-Multimer perform much better than other baselines on all the metrics. Specifically, xTrimoDock and AlphaFold-Multimer achieve a perfect MHQ score (100%), however, xTrimoDock’s RMSD, TM-Score and DockQ are better than AlphaFold-Multimer.

**Figure 6:**
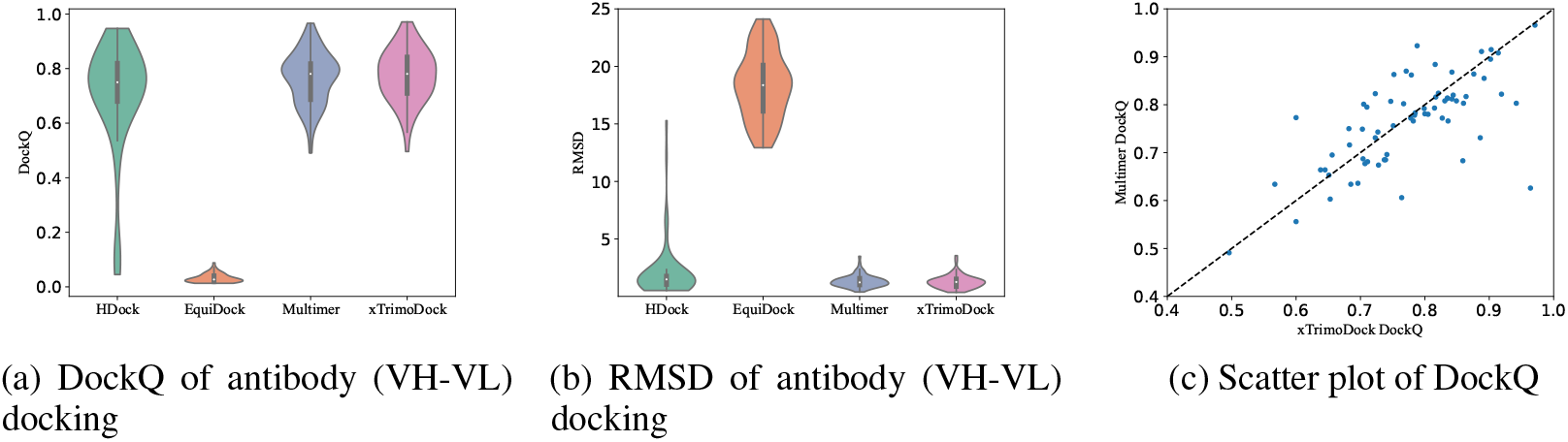
Result for antibody docking. Multimer refers to AlphaFold-Multimer.The larger DockQ indicates better results while the smaller RMSD indicates better results. EquiDock fails in most cases, while xTrimoDock has smaller RMSD and larger DockQ than AlphaFold-Multimer.

Typically, the CDRs on antibody are much more important than other regions because they contain binding sites in antibody-antigen docking. Because of the high variability and diverse conformations, it’s difficult to predict structures for CDRs. Table 2 provides a further comparison of xTrimoDock and AlphaFold-Multimer by comparing RMSD metrics on CDRs. The experiments show that xTrimoDock outperforms AlphaFold-Multimer on CDRs, especially on VH-CDR3 and VL-CDR3.

**Table 2:** The RMSD metrics of CDRs domain. It shows RMSD on CDR1, CDR2, CDR3 domain of both VH and VL respectively. xTrimoDock yields the best performance on CDR3 loop, which is a difficult domain to predict.

|  |  | AlphaFold-Multimer | xTrimoDock |
| --- | --- | --- | --- |
| VH | CDR1 | 1.232±1.104 | **1.205±1.009** |
|  | CDR2 | **0.938±0.616** | 0.973±0.580 |
|  | CDR3 | 2.755±1.500 | **2.644±1.697** |
| VL | CDR1 | 0.856±0.664 | **0.843±0.672** |
|  | CDR2 | **0.719±0.701** | 0.740±0.774 |
|  | CDR3 | 1.133±0.733 | **1.051±0.667** |

#### Antibody-antigen Docking

The results of antibody-antigen docking are summarized in Table 3 and Figure 7. Our xTrimoDock perform much better on the most important two metrics: DockQ and MHQ, than other baselines (10% improvement on DockQ, 16% improvement on MHQ compared to AlphaFold-Multimer). The scatter plot in Figure 7c illustrates a higher concentration of cases below the diagonal, indicating that xTrimoDock produces more accurate antibody-antigen complexes than AlphaFold-Multimer. The classical docking algorithms including ZDock, ClusPro and HDock as well as the deep learning method EquiDock fail in the most antibody-antigen docking cases.

**Table 3:** Experimental results of antibody and antibody-antigen docking. Antibody docking uses a light chain and heavy chain as input to assemble the antibody and the antibody-antigen docking takes the output of antibody docking and a predicted antigen structure as input. xTrimoDock outperforms other baselines in RMSD, TM-Score, DockQ and MHQ. *xTrimoDock w/o ML* refers to ablation study setting where distance prediction model is trained without masked MSA loss.

|  | Antibody VH-VL Docking |  |  |  | Antibody-Antigen Docking |  |  |  |
| --- | --- | --- | --- | --- | --- | --- | --- | --- |
|  | RMSD ↓ | TM-Score ↑ | DockQ ↑ | MHQ ↑ | RMSD ↓ | TM-Score ↑ | DockQ ↑ | MHQ ↑ |
| **ZDock** | 10.982±3.864 | 0.596±0.075 | 0.108±0.134 | 5.9% | 18.892±3.757 | 0.504±0.059 | 0.035±0.031 | 0.0% |
| **ClusPro** | 5.899±5.688 | 0.792±0.156 | 0.404±0.277 | 42.6% | 19.382±7.110 | 0.439±0.222 | 0.092±0.178 | 4.4% |
| **EquiDock** | 18.293±2.871 | 0.559±0.017 | 0.032±0.016 | 0.0% | 18.468±2.706 | 0.502±0.096 | 0.043±0.017 | 0.0% |
| **HDock** | 2.032±2.388 | 0.926±0.100 | 0.705±0.201 | 89.7 % | 15.779±6.364 | 0.612±0.137 | 0.090±0.187 | 7.4% |
| **AlphaFold-Multimer** | 1.325±0.530 | 0.962±0.020 | 0.765±0.094 | 100.0% | 13.650±5.886 | 0.640±0.124 | 0.108±0.172 | 7.4% |
| **xTrimoDock w/o ML** | 1.271±0.586 | 0.964±0.021 | 0.763±0.100 | 98.5% | 10.398±7.748 | 0.729±0.174 | 0.215±0.230 | 20.6% |
| **xTrimoDock** | **1.264±0.572** | **0.965±0.097** | **0.775±0.021** | **100.0%** | **10.090±7.817** | **0.733±0.173** | **0.220±0.232** | **23.5%** |

**Figure 7:**
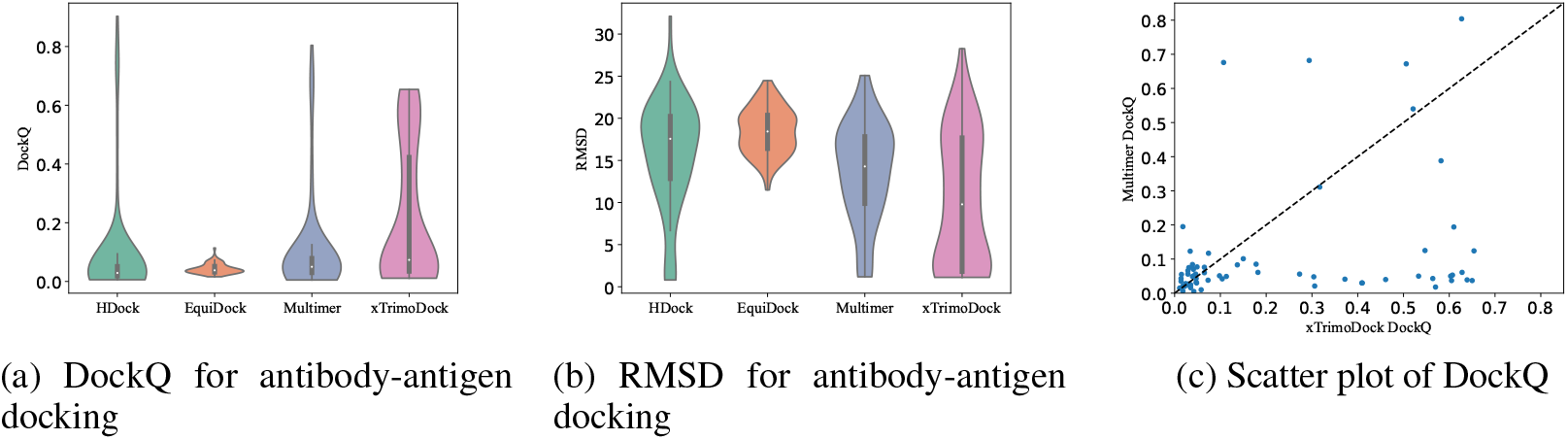
Result for antibody-antigen docking. Multimer refers to AlphaFold-Multimer. (a) and (b) show violin plots of DockQ and RMSD for antibody-antigen docking. More cases distribute below the diagonal in (c), indicating that xTrimoDock predicts better structures in more cases than AlphaFold-Multimer does.

#### Effects of Masked MSA Loss

We conduct an ablation study on xTrimoDock to investigate the effectiveness of masked MSA loss. We remove the MSA head from distance map prediction model in the training process. Table 3 shows a slight decrease in the metrics of both antibody docking and antibody-antigen docking. The MHQ of antibody docking decreases from 100.0% to 98.5% and the MHQ of antibody-antigen docking decreases from 23.5% to 20.6%. This suggests that MSA loss plays a key role in improving distance map prediction and can further affect the performance of docking. Masked MSA loss learns coevolutionary information from the paired and clustered MSAs and helps distance map learn better representation over cross chains.

### 4.3 Speed

The computational time for 68 cases in displayed in Table 4. As expected, traditional docking methods have longer inference times due to the intensive sampling, scoring, and refinement processes. AlphaFold-Multimer consumes a significant amount of time in searching MSA, taking approximately 1.5 hours per sample. xTrimoDock employs the accelerated MSA search algorithm MMseqs2 (Steinegger and Söding, 2017), and utilizes HPC optimizations such as layer norm (Cheng et al., 2022) (See Appendix C for details), resulting in a speedup of our algorithm by about 61 times compared to the original AlphaFold-Multimer. An ablation study removing spectral estimation shows that spectral estimation can shorten inference time significantly from 1.64 hours to 1.12 hours, suggesting that spectral estimation provides a good initialization for gradient refinement (see details in section 3.3 and section 3.4).

**Table 4:** Inference time of different methods on antibody-antigen docking. (unit:hour). Ablation study setting: *xTrimoDock w/o SE* refers to ablation study setting where xTrimoDock removes spectral estimation in section 3.3.

| Methods | Inference time |
| --- | --- |
| **ZDock** | 72.08 |
| **ClusPro** | 87.04 |
| **HDOCK** | 4.30 |
| **EquiDock** | 0.09 |
| **AlphaFold-Multimer** | 100.03 |
| **xTrimoDock w/o SE** | 1.64 |
| **xTrimoDock** | 1.12 |

Compared to classical docking softwares such ZDock and ClusPro, deep learning methods speed up the process by 10 to 150 times, which makes a huge difference in drug design. For instance, the human proteome contains up to 100,000 protein types, and a novel drug might negatively inhibit essential proteins. Therefore, the current hope is to scan for these interactions in a computational manner before bringing a few promising candidates to in vitro and in vivo testing.

### 4.4 Visualization

To more clearly and intuitively explore the advantages and weaknesses of xTrimoDock, we select three samples for visualization and analysis according to the type of antigen. We select two spike glycoproteins with *pdb*_*id* = 8*DLS* and *pdb*_*id* = 7*F* 6*Z*, one circumsporozoite protein with *pdb*_*id* = 7*RXI*. The ground truth structures and predictions of competitive methods are shown in Figure 8. It can be demonstrated that xTrimoDock is able to provide competitive docking pose on antibody-antigen complexes.

**Figure 8:**
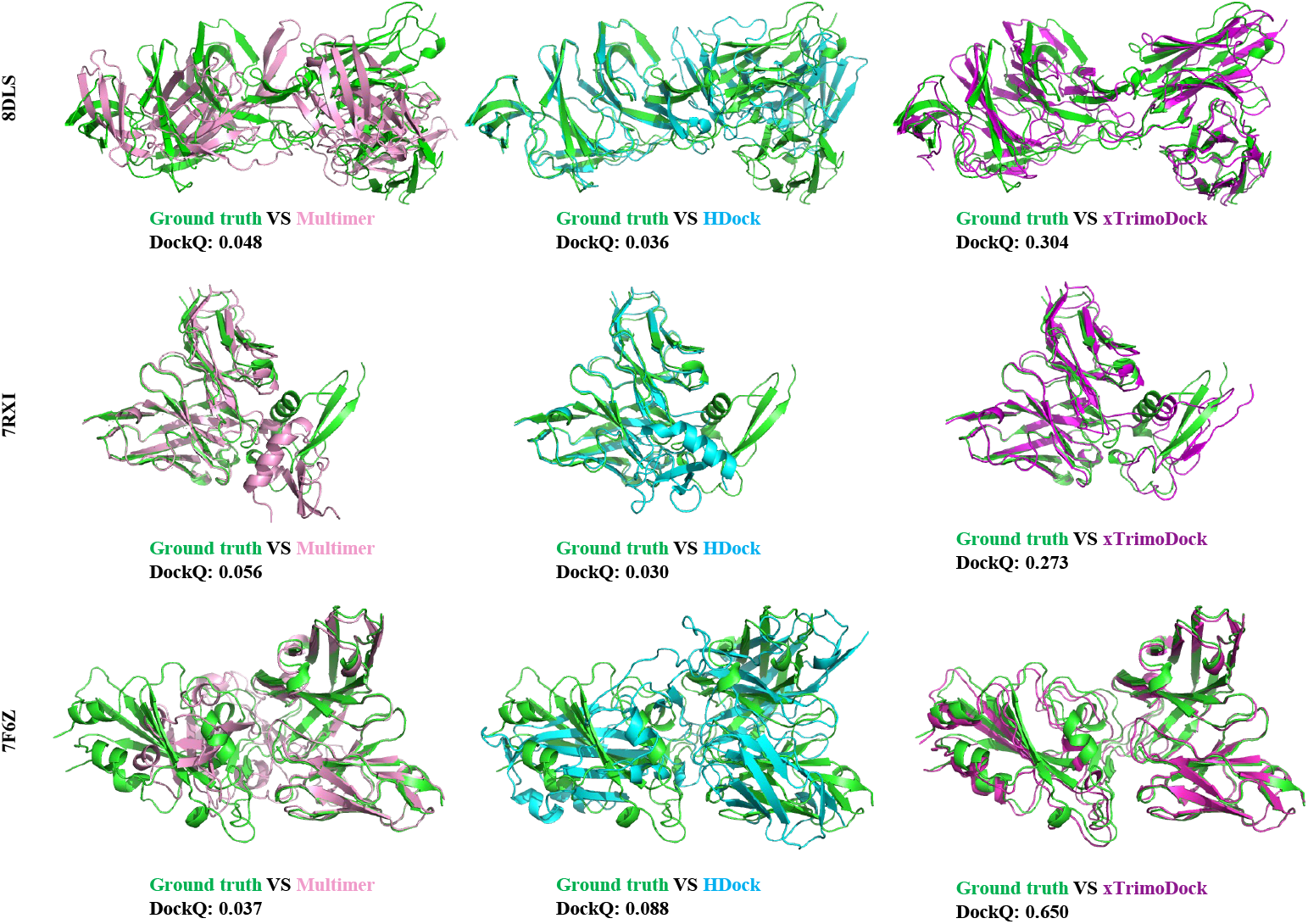
Structure comparison between the predicted structures with the ground truth structure of 8DLS, 7RXI and 7F6Z. The structures of the ground truth, Multimer, HDock and xTrimoDock are colored green, pink, cyan and magenta, respectively. The type of antigens in 8DLS and 7F6Z belongs to spike glycoprotein and the type of antigen in 7RXI belongs to circumsporozoite.

## 5 Conclusion

In this paper, we have presented a novel multi-chain rigid docking approach that utilizes crossmodal representation learning and spectral algorithm to obtain rotation-translation transformation for docking. As far as we know, it is first deep learning approach which uses a contact map and rigid docking physics as the key component for multi-chain docking. Moreover, the use of cross-modal representation learning to obtain the contact map enables us to circumvent the use of Multimer’s structure models in multi-chain prediction scenarios. Experiments on antibody and antibody-antigen docking demonstrate competitive results.

A limitation of xTrimoDock is its inability to consider protein flexibility, which can play a significant role in the real-world protein docking process. Changes in the structure of interacting proteins can be substantial, making it an area of future work to extend xTrimoDock to handle flexible docking. This will be a more challenging but highly valuable task in drug design. Finally, we hope that our work will bring greater attention to the use of deep learning in biological applications.

---

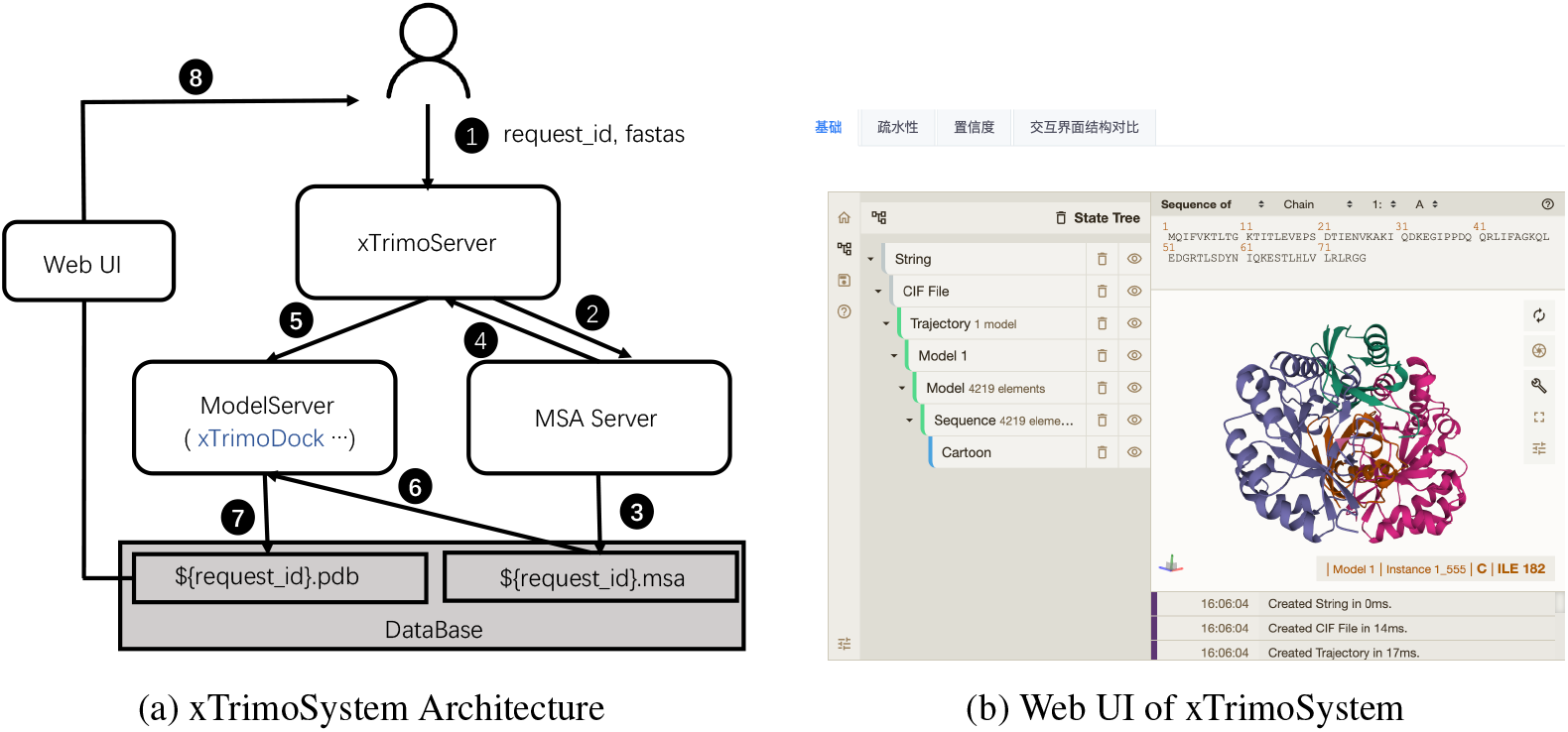

## A DataSet

We collected a dataset containing 68 samples and used this dataset to generate two test sets, the first antibody dataset and the second antibody-antigen dataset. These 68 samples are respectively 8dls, 8dlr, 8dfi, 8dfh, 8dcc, 8dad, 7zr8, 7zf8, 7xxl, 7xh8, 7×26, 7wsl, 7wsi, 7ws6, 7ws2, 7wrz, 7wrv, 7wro, 7wrl, 7wrj, 7wog, 7wlc, 7wef, 7wee, 7wed, 7wcr, 7wbz, 7urq, 7uaq, 7tty, 7ttx, 7ttm, 7tpj, 7tp4, 7tp3, 7tlz, 7the, 7tc9, 7t8w, 7t7b, 7t01, 7swp, 7su1, 7str, 7sem, 7sd5, 7sbu, 7sbg, 7sbd, 7sa6, 7s5p, 7rxp, 7rxi, 7rbu, 7qtk, 7n0a, 7lo8, 7lo7, 7kql, 7fjc, 7f7e, 7f6z, 7f6y, 7eng, 7ek0, 7ejz, 7ejy, 7e9p.

## B MSA Mask Policy

We randomly mask each position in a MSA cluster center with 15% probability. Each masked token in the MSA is replaced by other token with the following policy:

- 70% probability of substitution with a special token (token=⋆).
- 10% probability of substitution with a uniformly sampled random amino acid.
- 10% probability of substitution with an amino acid sampled from the Multiple Sequence Alignment (MSA) profile for the corresponding position.
- 10% probability of no substitution.

## C Deployment on xTrimoSystem

xTrimoSystem is an advanced high-performance protein structure processing architecture designed to decouple Multi Sequence Alignments (MSA) and neural network computation (e.g., Evoformer and Structure Prediction). The system is comprised of three key components: xTrimoServer, MSA retrieval server, and ModelServer (a neural network inference server).

xTrimoServer offers efficient streaming dispatch and reduces the dependence of neural network inference on MSA, thereby maximizing server overlap and optimizing resource utilization. As a result, the average protein prediction time is reduced from approximately 800s to just 40s, representing a 20-fold acceleration. For instance, xTrimoServer consistently receives and sends samples (e.g., request_id and fasta) to the MSA server, which processes a single MSA with maximum concurrency, writes the MSA to the database in a timely manner, and feeds the result back to the inference server. This allows xTrimoServer to promptly start processing the next request. Once the request is sent to ModelServer, it retrieves the corresponding MSA from the database using the request_id and infers the protein structure (saved as pdb files in the database). xTrimoDock is deployed in the ModelServer.

The front-end of the interface with xTrimoSystem is designed for 3D vision and protein structure analysis and is an extension of the molstar web app plugin (https://github.com/molstar/molstar). The Web UI has been inherited in our system to provide an intuitive and user-friendly interface.

## D Further Introduction on Signatures and Model Inputs

There contains three types of signatures: structure signatures, MSA signatures and sequence signatures.

### Structure Signatures

We derive two types of features from single-chain structures: *rigid angle feature* 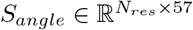 and *rigid pair feature* 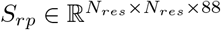.

- The *rigid angle feature* contains a) one-hot representation of primary sequences of size [*N*_*res*_, 22] (20 amino acids, 1 unknown type and 1 gap or missing residue), b) angle features of size [*N*_*res*_, 28] (sine and cosine representation to encode three backbone torsion angles and four side-chain torsion angles as well as alternative torsion angles with 180^*o*^-rotation symmetry for each residue’s local frame), c) angle indicator feature of size [*N*_*res*_, 7] (indicating whether the torsion angles exist).
- The *rigid pair feature* contains: a) distogram feature of size [*N*_*res*_, *N*_*res*_, 39] (discretized distance between beta carbons atoms, alpha carbon used for glycine in intra-chain and inter-chain. The pairwise distances are discretized into 38 bins of equal width between 3.25 Å and 50.75 Å and one more bin contains any larger distances), b) residue type feature of size [*N*_*res*_, *N*_*res*_, 44] (expanded from residue one-hot representation of size [*N*_*res*_, 1, 22] and [*N*_*res*_, 22, 1]), c) backbone feature of size [*N*_*res*_, *N*_*res*_, 3] (build local frame coordinates using Gram–Schmidt process based on the original *N, Cα, C* coordinates, extract the unit vector as backbone feature), d) residue indicator feature of size [*N*_*res*_, *N*_*res*_, 1] (expanded from residue existing indicator [1, *N*_*res*_, 1]), e) pair indicator feature of size [*N*_*res*_, *N*_*res*_, 1] (indicating whether pair exists).

### MSA Signatures

Cross-chain coevolutionary information is a useful feature in distance predictions. We employ the heuristic approach in (Evans et al., 2021) to pair sequences between per-chain MSAs. Specifically, the per-chain MSA sequences are grouped by species, where the species labels derived from UniProt’s idmapping ^7^. Then the grouped sequences are paired within a specific species. We concatenate the sequences if there is only one sequence per species. Otherwise, we match the chain MSAs by minimizing the base-pair distance between chains for prokaryotic species and match the chain MSAs by ordering them by sequence identity to the target sequence for eukaryotic species (Zhou et al., 2018). To reduce the computational and memory cost, we employ the MSA clustering approach in AlphaFold2 (Jumper et al., 2021). Specifically, we select *N*_*cluster*_=252 sequences randomly as MSA cluster centers, where we set the first cluster center always be the primary protein sequence. The remaining sequences are assigned to their closest cluster by Hamming distance. Also, we select *N*_*extra*_=1024 sequences randomly without replacement from sequences that have not been selected as cluster centers. Finally, we derive two types of MSA features: *cluster MSA feature* 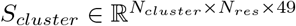 and *extra MSA feature* 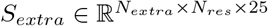.

- *cluster MSA feature* contains a) one-hot representation of size [*N*_*cluster*_, *N*_*res*_, 23] (20 amino acids, 1 unknown type, 1 gap or missing residue, and a mask token which will be introduced in section 3.2.3), b) amino acid distribution of size [*N*_*cluster*_, *N*_*res*_, 23] (the distribution over amino acid types in each MSA cluster, where) c) deletion indicator of size [*N*_*cluster*_, *N*_*res*_, 1] (whether there is a deletion to the left of the residue) d) deletion value of size [*N*_*cluster*_, *N*_*res*_, 1] (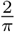 arctan 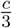, where *c* refers to the number of deletion of the left of every position) e) the mean deletion value of size [*N*_*cluster*_, *N*_*res*_, 1] (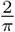 arctan 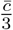, where 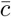 refers to the average number of deletion of the left of every position)
- We derive the similar features a), c) and d) from extra MSAs.

### Sequence Signatures

We derive two types of feature from primary sequence: *residue feature* 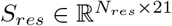 and *primary pair feature* 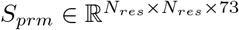.

- The *residue feature* contains an one-hot feature of size [*N*_*res*_, 21], including 20 amino acids and 1 unknown type.
- In general, we add three chain identifiers, *entity_id, asym_id, sym_id*, for each residue, where *entity_id* distinguishes unique chain sequences, *asym_id* distinguishes between chains, *sym_id* distinguishes chains who have the same sequence ^8^. To provide the network with amino acids position information within or cross chains, we also derive *primary pair feature* of size [*N*_*res*_, *N*_*res*_, 73], which contains: a) relative positional feature of size [*N*_*res*_, *N*_*res*_, 66] (relative residue indices are clipped between [-32, 32]. We assign the cross-chain pair by the 66-th index.), b) entity indicator of of size [*N*_*res*_, *N*_*res*_, 1] (whether residues *i* and *j* in pair *ij* come from the same chain), c) relative index feature of size [*N*_*res*_, *N*_*res*_, 6] (relative *sym_id* indices are clipped between [-2, 2]. We assign the pair whose two residues have different *sym_ids* by 6-th index.)

## Footnotes

3 https://ftp.uniprot.org/pub/databases/uniprot/current_release/knowledgebase/idmapping/

4 https://github.com/aqlaboratory/openfold

5 https://github.com/octavian-ganea/equidock_public

6 https://github.com/bjornwallner/DockQ

7 https://ftp.uniprot.org/pub/databases/uniprot/current_release/knowledgebase/idmapping/

8 In general, our contact prediction model can solve cases more than three chains. We provide an example to introduce *entity_id, asym_id, sym_id*. If complex contains fives chains: A, B, B, C, C, where A, B, C refers to three unique chains. The *entity_ids* of each chain are 1, 2, 2, 3, 3 The *sym_ids* of each chain are is 1, 1, 2, 1, 2. The *asym_ids* of each chain are is 1, 2, 3, 4, 5.

## References

Brennan Abanades, Guy Georges, Alexander Bujotzek, and Charlotte M Deane. 2022a. ABlooper: Fast accurate antibody CDR loop structure prediction with accuracy estimation. Bioinformatics 38, 7 (2022), 1877–1880.

Brennan Abanades, Wing Ki Wong, Fergus Boyles, Guy Georges, Alexander Bujotzek, and Charlotte M Deane. 2022b. ImmuneBuilder: Deep-Learning models for predicting the structures of immune proteins. bioRxiv (2022), 2022–11.

Gustaf Ahdritz, Nazim Bouatta, Sachin Kadyan, Qinghui Xia, William Gerecke, Timothy J O’Donnell, Daniel Berenberg, Ian Fisk, Niccolò Zanichelli, Bo Zhang, et al. 2022. OpenFold: Retraining AlphaFold2 yields new insights into its learning mechanisms and capacity for generalization. bioRxiv (2022), 2022–11.

Minkyung Baek, Frank DiMaio, Ivan Anishchenko, Justas Dauparas, Sergey Ovchinnikov, Gyu Rie Lee, Jue Wang, Qian Cong, Lisa N Kinch, R Dustin Schaeffer, et al. 2021. Accurate prediction of protein structures and interactions using a three-track neural network. Science 373, 6557 (2021), 871–876.

Itzhack Y Bar-Itzhack. 2000. New method for extracting the quaternion from a rotation matrix. Journal of guidance, control, and dynamics 23, 6 (2000), 1085–1087.

Sankar Basu and Björn Wallner. 2016. DockQ: a quality measure for protein-protein docking models. PloS one 11, 8 (2016), e0161879.

Helen M Berman. 2008. The protein data bank: a historical perspective. Acta Crystallographica Section A 64, 1 (2008), 88–95.

Jacek Biesiada, Aleksey Porollo, Prakash Velayutham, Michal Kouril, and Jaroslaw Meller. 2011. Survey of public domain software for docking simulations and virtual screening. Human genomics 5, 5 (2011), 1–9.

Rong Chen, Li Li, and Zhiping Weng. 2003. ZDOCK: an initial-stage protein-docking algorithm. Proteins: Structure, Function, and Bioinformatics 52, 1 (2003), 80–87.

Shenggan Cheng, Ruidong Wu, Zhongming Yu, Binrui Li, Xiwen Zhang, Jian Peng, and Yang You. 2022. FastFold: reducing AlphaFold training time from 11 days to 67 hours. arXiv preprint arXiv:2203.00854 (2022).

Charles Christoffer, Siyang Chen, Vijay Bharadwaj, Tunde Aderinwale, Vidhur Kumar, Matin Hormati, and Daisuke Kihara. 2021. LZerD webserver for pairwise and multiple protein–protein docking. Nucleic Acids Research 49, W1 (2021), W359–W365.

Sjoerd J De Vries, Marc Van Dijk, and Alexandre MJJ Bonvin. 2010. The HADDOCK web server for data-driven biomolecular docking. Nature protocols 5, 5 (2010), 883–897.

Stephan Eismann, Raphael JL Townshend, Nathaniel Thomas, Milind Jagota, Bowen Jing, and Ron O Dror. 2021. Hierarchical, rotation-equivariant neural networks to select structural models of protein complexes. Proteins: Structure, Function, and Bioinformatics 89, 5 (2021), 493–501.

Richard Evans, Michael O’Neill, Alexander Pritzel, Natasha Antropova, Andrew W Senior, Timothy Green, Augustin Žídek, Russell Bates, Sam Blackwell, Jason Yim, et al. 2021. Protein complex prediction with AlphaFold-Multimer. BioRxiv (2021).

Xiaomin Fang, Fan Wang, Lihang Liu, Jingzhou He, Dayong Lin, Yingfei Xiang, Xiaonan Zhang, Hua Wu, Hui Li, and Le Song. 2022. Helixfold-single: Msa-free protein structure prediction by using protein language model as an alternative. arXiv preprint arXiv:2207.13921 (2022).

Octavian-Eugen Ganea, Xinyuan Huang, Charlotte Bunne, Yatao Bian, Regina Barzilay, Tommi S. Jaakkola, and Andreas Krause. 2022. Independent SE(3)-Equivariant Models for End-to-End Rigid Protein Docking. In ICLR. OpenReview.net.

Octavian-Eugen Ganea, Xinyuan Huang, Charlotte Bunne, Yatao Bian, Regina Barzilay, Tommi Jaakkola, and Andreas Krause. 2021. Independent se (3)-equivariant models for end-to-end rigid protein docking. arXiv preprint arXiv:2111.07786 (2021).

L Steven Johnson, Sean R Eddy, and Elon Portugaly. 2010. Hidden Markov model speed heuristic and iterative HMM search procedure. BMC bioinformatics 11, 1 (2010), 1–8.

John Jumper, Richard Evans, Alexander Pritzel, Tim Green, Michael Figurnov, Olaf Ronneberger, Kathryn Tunyasuvunakool, Russ Bates, Augustin Žídek, Anna Potapenko, et al. 2021. Highly accurate protein structure prediction with AlphaFold. Nature 596, 7873 (2021), 583–589.

Dima Kozakov, David R Hall, Bing Xia, Kathryn A Porter, Dzmitry Padhorny, Christine Yueh, Dmitri Beglov, and Sandor Vajda. 2017. The ClusPro web server for protein–protein docking. Nature protocols 12, 2 (2017), 255–278.

Elodie Laine, Stephan Eismann, Arne Elofsson, and Sergei Grudinin. 2021. Protein sequence-to-structure learning: Is this the end (-to-end revolution)? Proteins: Structure, Function, and Bioinformatics 89, 12 (2021), 1770–1786.

Guillaume Launay, Masahito Ohue, Julia Prieto Santero, Yuri Matsuzaki, Cécile Hilpert, Nobuyuki Uchikoga, Takanori Hayashi, and Juliette Martin. 2020. Evaluation of consrank-like scoring functions for rescoring ensembles of protein–protein docking poses. Frontiers in molecular biosciences 7 (2020), 559005.

Zeming Lin, Halil Akin, Roshan Rao, Brian Hie, Zhongkai Zhu, Wenting Lu, Nikita Smetanin, Robert Verkuil, Ori Kabeli, Yaniv Shmueli, et al. 2022. Evolutionary-scale prediction of atomic level protein structure with a language model. bioRxiv (2022), 2022–07.

Michael Remmert, Andreas Biegert, Andreas Hauser, and Johannes Söding. 2012. HHblits: lightning-fast iterative protein sequence searching by HMM-HMM alignment. Nature methods 9, 2 (2012), 173–175.

Jeffrey A Ruffolo and Jeffrey J Gray. 2022. Fast, accurate antibody structure prediction from deep learning on massive set of natural antibodies. Biophysical Journal 121, 3 (2022), 155a–156a.

Jeffrey A Ruffolo, Jeremias Sulam, and Jeffrey J Gray. 2022. Antibody structure prediction using interpretable deep learning. Patterns 3, 2 (2022), 100406.

Christina EM Schindler, Isaure Chauvot de Beauchêne, Sjoerd J de Vries, and Martin Zacharias. 2017. Protein-protein and peptide-protein docking and refinement using ATTRACT in CAPRI. Proteins: Structure, Function, and Bioinformatics 85, 3 (2017), 391–398.

Andrew W Senior, Richard Evans, John Jumper, James Kirkpatrick, Laurent Sifre, Tim Green, Chongli Qin, Augustin Žídek, Alexander WR Nelson, Alex Bridgland, et al. 2019. Protein structure prediction using multiple deep neural networks in the 13th Critical Assessment of Protein Structure Prediction (CASP13). Proteins: Structure, Function, and Bioinformatics 87, 12 (2019), 1141–1148.

Martin Steinegger and Johannes Söding. 2017. MMseqs2 enables sensitive protein sequence searching for the analysis of massive data sets. Nature biotechnology 35, 11 (2017), 1026–1028.

Sharon Sunny and PB Jayaraj. 2021. FPDock: Protein–protein docking using flower pollination algorithm. Computational Biology and Chemistry 93 (2021), 107518.

Mieczyslaw Torchala, Iain H Moal, Raphael AG Chaleil, Juan Fernandez-Recio, and Paul A Bates. 2013. SwarmDock: a server for flexible protein–protein docking. Bioinformatics 29, 6 (2013), 807–809.

Ilya A Vakser. 2014. Protein-protein docking: From interaction to interactome. Biophysical journal 107, 8 (2014), 1785–1793.

Vishwesh Venkatraman, Yifeng D Yang, Lee Sael, and Daisuke Kihara. 2009. Protein-protein docking using region-based 3D Zernike descriptors. BMC bioinformatics 10, 1 (2009), 1–21.

Jacob Verburgt and Daisuke Kihara. 2022. Benchmarking of structure refinement methods for protein complex models. Proteins: Structure, Function, and Bioinformatics 90, 1 (2022), 83–95.

Yining Wang, Xumeng Gong, Shaochuan Li, Bing Yang, YiWu Sun, Chuan Shi, Hui Li, Yangang Wang, Cheng Yang, and Le Song. 2022. xTrimoABFold: Improving Antibody Structure Prediction without Multiple Sequence Alignments. arXiv preprint arXiv:2212.00735 (2022).

Gaoqi Weng, Ercheng Wang, Zhe Wang, Hui Liu, Feng Zhu, Dan Li, and Tingjun Hou. 2019. Hawk-Dock: a web server to predict and analyze the protein–protein complex based on computational docking and MM/GBSA. Nucleic acids research 47, W1 (2019), W322–W330.

Ruidong Wu, Fan Ding, Rui Wang, Rui Shen, Xiwen Zhang, Shitong Luo, Chenpeng Su, Zuofan Wu, Qi Xie, Bonnie Berger, et al. 2022. High-resolution de novo structure prediction from primary sequence. BioRxiv (2022), 2022–07.

Sitao Wu and Yang Zhang. 2008. A comprehensive assessment of sequence-based and template-based methods for protein contact prediction. Bioinformatics 24, 7 (2008), 924–931.

Yumeng Yan, Huanyu Tao, Jiahua He, and Sheng-You Huang. 2020. The HDOCK server for integrated protein–protein docking. Nature protocols 15, 5 (2020), 1829–1852.

Chengxin Zhang, Morgan Shine, Anna Marie Pyle, and Yang Zhang. 2022. US-align: Universal Structure Alignments of Proteins, Nucleic Acids, and Macromolecular Complexes. bioRxiv (2022).

Yang Zhang and Jeffrey Skolnick. 2004. Scoring function for automated assessment of protein structure template quality. Proteins: Structure, Function, and Bioinformatics 57, 4 (2004), 702– 710.

Tian-ming Zhou, Sheng Wang, and Jinbo Xu. 2018. Deep learning reveals many more inter-protein residue-residue contacts than direct coupling analysis. BioRxiv (2018), 240754.

